# Two-color nanoscopy of organelles for extended times with HIDE probes

**DOI:** 10.1101/647065

**Authors:** Ling Chu, Jonathan Tyson, Juliana E. Shaw, Felix Rivera-Molina, Anthony J. Koleske, Alanna Schepartz, Derek K. Toomre

## Abstract

Performing multi-color nanoscopy for extended times is challenging due to the rapid photobleaching rate of most fluorophores. Here we describe a new fluorophore (Yale-595) and a bio-orthogonal labeling strategy that enables both super-resolution (STED) and 3D confocal imaging of two organelles simultaneously for extended times using high-density environmentally sensitive (HIDE) probes. Because HIDE probes are small, cell-permeant molecules, they can visualize organelle pairs (ER + mitochondria, ER + plasma membrane) in hard-to-transfect cell lines at super-resolution for up to 7 minutes. The extended time domain possible using these new tools reveal novel dynamic nanoscale targeting between organelles.

---

Super-resolution microscopy (‘nanoscopy’) can visualize cellular components with resolutions as high as ~10 nanometers, far below the Abbe diffraction limit^1^. When combined with multi-color labeling strategies, nanoscopy can reveal key features of organelle structure and interactions that remain opaque when visualized using diffraction-limited approaches^2, 3^. However, visualizing organelle *dynamics* at super-resolution remains challenging because even the most photostable fluorophores bleach within tens to hundreds of seconds under conditions required for STED or SMS nanoscopy^4^.

Recently, we reported a set of high-density environmentally sensitive (HIDE) probes that support long time-lapse, single-color nanoscopy of organelles including the ER, mitochondria, Golgi apparatus, and plasma membrane (PM)^5–7^. HIDE probes consist of an organelle-specific lipid or lipid-like small molecule with a reactive *trans*-cyclooctene (TCO) moiety and a silicon rhodamine-tetrazine (SiR-Tz)^8^ reaction partner. These two components undergo a rapid, *in situ* tetrazine ligation reaction^9, 10^ that localizes the SiR dye at high density within the organelle membrane. In this environment, HIDE probes support the acquisition of continuous SMS and STED super-resolution movies in standard culture media (no redox chemicals) for up to 25 mins, approximately 50-fold longer than when the identical fluorophore is linked to an organelle-resident protein^7^. Here we describe innovations that enable live-cell HIDE imaging of two different organelles in two colors for extended times using STED (Figure 1a). The ~50 nm per axis resolution achieved here significantly exceeds the ~100 nm resolution obtained when using structured illumination^11^ or lattice light-sheet microscopy^12^.

**Figure 1.**
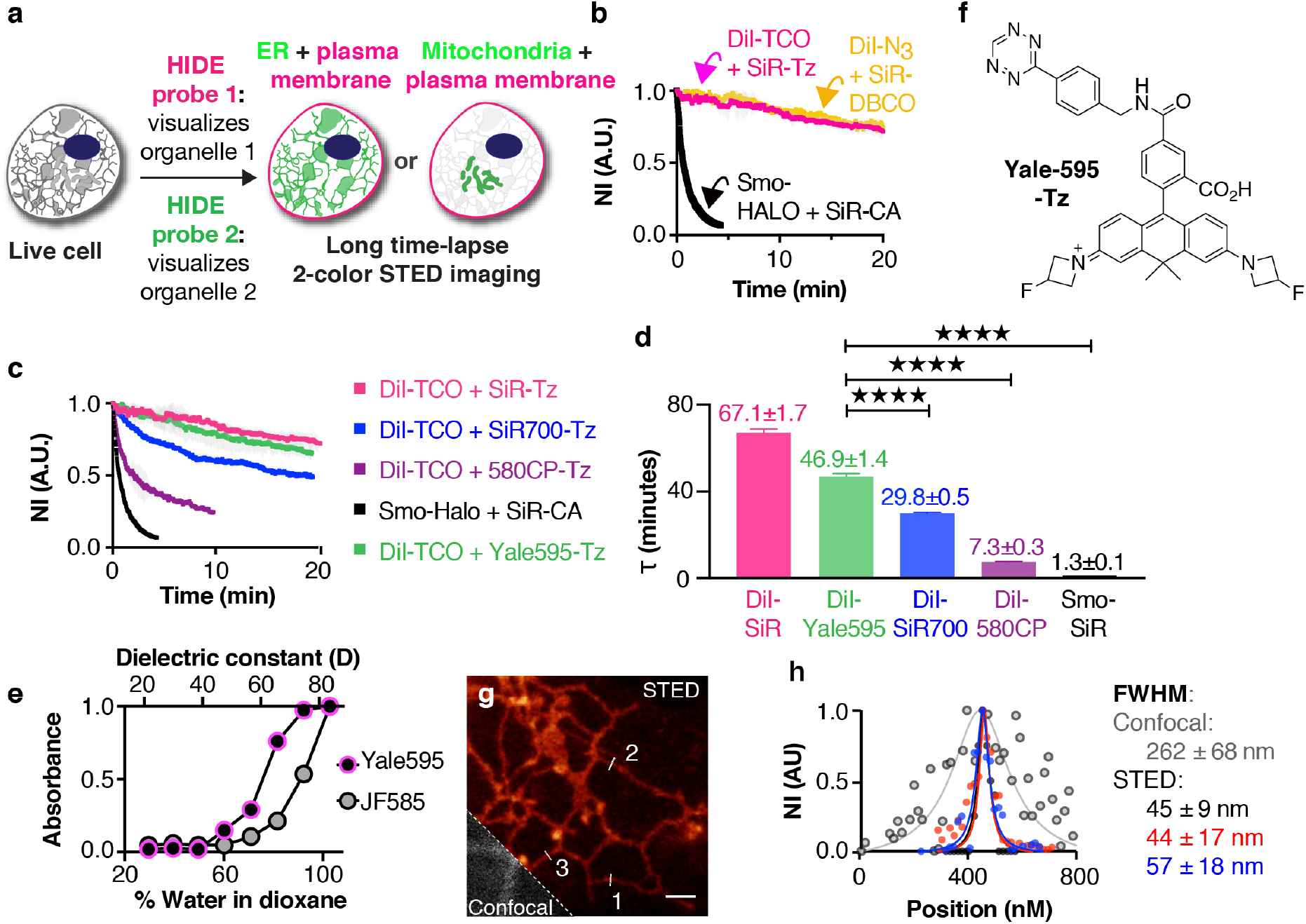
Development of two-color high-density environment-sensitive (HIDE) probes. (**a**) HIDE probes enable long time-lapse two-color STED imaging. (**b**) Plot illustrating normalized fluorescence intensity of Smo-HALO+SiR-CA (blue), Dil-TCO+SiR-Tz (red) and Dil-N_3_+SiR-DBCO (green) over time (mean±SD, n = 3 ROI, N = 1 cell). (**c**) Plot illustrating normalized fluorescence intensity (NI) of 580CP (purple), SiR (red), SiR700 (blue) and Yale595 (green) over time (mean±SD, n = 3 ROI, N = 1 cell). (**d**) τ values calculated from a single exponential fit to the photobleaching curves in **c** (mean ± SD., n = 3 ROI, N = 1 cell). ****P ≤ 0.0001. (**e**) Plot of normalized absorbance of Yale595-COOH and JF585-COOH in response to different dielectric constant, D, of dioxane-water mixtures (mean, n = 2). (**f**) Chemical structure of Yale595-Tz. (**g**) STED imaging of endoplasmic reticulum of HeLa cells labeled with 2 μM Cer-TCO for 1 hr followed by 2 μM Yale595-Tz for 30 min with a confocal cutaway in gray. Scale bar: 1 μm (**h**) A plot of the fluorescence signal in the STED and confocal image as a function of position along the line profile in **g.** A fit of three line profiles from the confocal data to a Lorentzian function (grey line) provides a FWHM (mean ± SD, n = 3) of 262 ±68 nm. Fits of three line profiles (white lines) in the STED image individually to a Lorentzian function provide FWHM values of 45 ±9 nm, 44 ± 17 nm, and 57 ± 18 nm.

Two-color, live-cell HIDE nanoscopy demands not only two organelle-specific small molecules and a second photostable fluorophore but also a second conjugation reaction that is orthogonal to the tetrazine ligation and proceeds rapidly to completion within live cells. We were drawn to the strain-promoted azide-alkyne cycloaddition reaction (SPAAC)^13^ between an azide and a dibenzoazacyclooctyne (DBCO) as it is live-cell compatible and rapid enough (*k* = 0.31 M^−1^s^−1^)^14^ to ensure complete reaction using reagents at micromolar concentrations.

To confirm that a SPAAC-assembled HIDE probe would localize correctly, DiI-N_3_ and SiR-DBCO were synthesized (Supplementary Information, Figure S1a), added sequentially to HeLa cells, incubated 30 minutes to generate DiI∗SiR and imaged by confocal microscopy (Figure S2). Under these conditions, DiI∗SiR colocalized extensively with a bona fide PM marker, VAMP2-pH (PCC = 0.63 ± 0.06), no colocalization was observed when cells were incubated with SiR-DBCO but not DiI-N_3_ (PCC = 0.29 ± 0.03) (Figure S2c). As observed previously^5^, the HIDE probe DiI∗SiR must be assembled in two steps, as cells treated with the pre-assembled reaction product DiI∗SiR (Figure S1a) showed no PM labeling in SiR channel (Figure S3).

To test whether a HIDE probe assembled using SPAAC would support prolonged STED nanoscopy, we treated HeLa cells, under optimized conditions (Figure S4), with 10 μM DiI-N_3_ and 2 μM SiR-DBCO. We imaged the cells continuously by STED for 20 minutes at 0.5 Hz and monitored plasma membrane (PM) fluorescence over time (Movie S1). Both the initial fluorescence intensity (Figure S5a) and lifetime (τ) (Figure 1b and S5b) of the HIDE probe generated from DiI-N_3_ and SiR-DBCO were virtually identical to those measured using DiI-TCO and SiR-Tz. The similarity of these values confirms that HIDE probes assembled using SPAAC support prolonged STED imaging; the small difference in structure due to alternative linker chemistry has no apparent effect on SiR photostability. In contrast, STED images generated using the Smo-Halo/SiR-CA combination bleached within 1 minute when visualized by STED (Movie S2 and Figure 1b).

Next, we sought to identify a STED-appropriate fluorophore that would be compatible with SiR to enable two-color HIDE nanoscopy. The ideal second fluorophore should be: (1) membrane-permeant; (2) non-toxic; (3) highly photostable; and (4) spectrally separable from SiR. We first evaluated four previously reported fluorophores: 580CP^15^, Atto590^16, 17^, JF585^18^, and SiR700^19^ (Figure S6). HeLa cells were treated with 10 μM DiI-TCO followed by 2 μM of either 580CP-Tz, Atto590-Tz, JF585-Tz, or SiR700-Tz (Supplementary Information) then imaged by STED at 0.5 Hz. None of these previously reported fluorophores were suitable. Visual inspection and time-dependent quantification of PM fluorescence revealed that the HIDE probes generated from DiI-TCO and either CP580-Tz or SiR700-Tz bleached considerably more rapidly than that generated using SiR-Tz (Figure 1c, Figure S7; Movies S2) and HIDE probe generated from DiI-TCO and SiR700-Tz was also dimmer (Figure S7d; Movies S3). The HIDE probe generated from DiI-TCO and Atto590-Tz was acutely cytotoxic (Movie S4), while that generated from DiI-TCO and JF585-Tz was extremely dim (Figure S7b and d). Notably, all PM-localized HIDE probes generated here could be imaged for an order of magnitude longer then a PM-localized protein Smo-Halo/SiR-CA (Movie S5).

We were intrigued by the poor performance of the HIDE probe generated from DiI-TCO and JF585-Tz, as this fluorophore performs well when used to label a membrane resident protein^18^. We hypothesized that this difference reflected a shift in the equilibrium between the open (ON) and closed (OFF) states of this dye in the two environments: JF-585 performs well when attached to a membrane protein because it is mainly ON in a polar, aqueous environment, but poorly as a HIDE probe because it is mainly OFF in the lipid environment. A plot of JF585 emission as a function of the percent water in a water/dioxane mixture is consistent with this hypothesis; the midpoint of the transition occurs at approximately 91% water (Figure 1e). We reasoned that an analog of JF585 in which the 3,3-difluoroazetidine ring is replaced with less electrophilic 3-fluoroazetidine (Supplementary Information, Yale595-Tz, Figure 1f) would display a higher ON fraction in the lipid environment and thereby facilitate STED imaging. The excitation and emission maxima of Yale595-CO_2_H were 597 nm and 621 nm, respectively, values that are compatible with SiR for two-color imaging (Figure S8a). As anticipated, a plot of Yale595 emission as a function of the percent water in a water/dioxane mixture was shifted, with a midpoint at 75% water (Figure 1e). Moreover, the apparent photostabilities of HIDE probes generated from DiI-TCO and either Yale595-Tz or SiR-Tz were similar (Figure 1c, d, Figure S7 and Movie S6). To further validate the utility of Yale595 in cells, we generated an ER-specific HIDE probe from Cer-TCO^5^ and Yale595-Tz. This probe colocalized with Sec61b-GFP (Figure S9c and d) and when visualized using STED could resolve individual ER tubules with a FWHM of ~50 nm (Figure 1g and h).

The ER HIDE probe generated using Yale595-Tz was then used in combination with DiI-N_3_/SiR-DBCO to achieve two-color HIDE imaging of the ER and PM. HeLa cells were treated first with DiI-N_3_ and Cer-TCO then with SiR-DBCO and Yale595-Tz (Figure 2a, Figure S10) and imaged by STED (Figure 2b). High resolution images were obtained in both channels (FWHM = 72 ± 19 nm for Yale 595; 57 ± 8 nm for SiR (Figure 2c) with cross-talk between the two channels minimized by spectral unmixing (Figure S11). With the two-color HIDE probes, we observed ER tubule formation, elongation, and rearrangement as well as filopodia movement over 250 time points with no significant loss of fluorescence (Figure 2d, e, Movie S7). When conjugated to ER (Sec61β)- and PM (Smo)-resident proteins through Halo and SNAP tags, respectively, SiR and Yale595 bleached 15- to 40-fold faster (Figure 2d, e, Movie S8). The two-color HIDE strategy was easily generalized to simultaneously image the PM and mitochondria in live cells for extended times. HeLa cells were treated with DiI-N_3_ along with the mitochondria-specific small molecule RhoB-TCO followed by SiR-DBCO and Yale595-Tz (Figure 2f, Figure S12) then imaged by STED at 0.5 Hz (Figure 2g, Movie S9). Here the fine and rapid dynamic processes of mitochondria fission and filopodia remodeling could be observed and studied over 5 minutes with minimal photobleaching. The toxicity of two-color HIDE probes was studied by monitoring cell division events of HeLa cells. No toxicity of the probes was observed, as monitored by the number of cell division events per hour after probe treatment (Figure S13).

**Figure 2.**
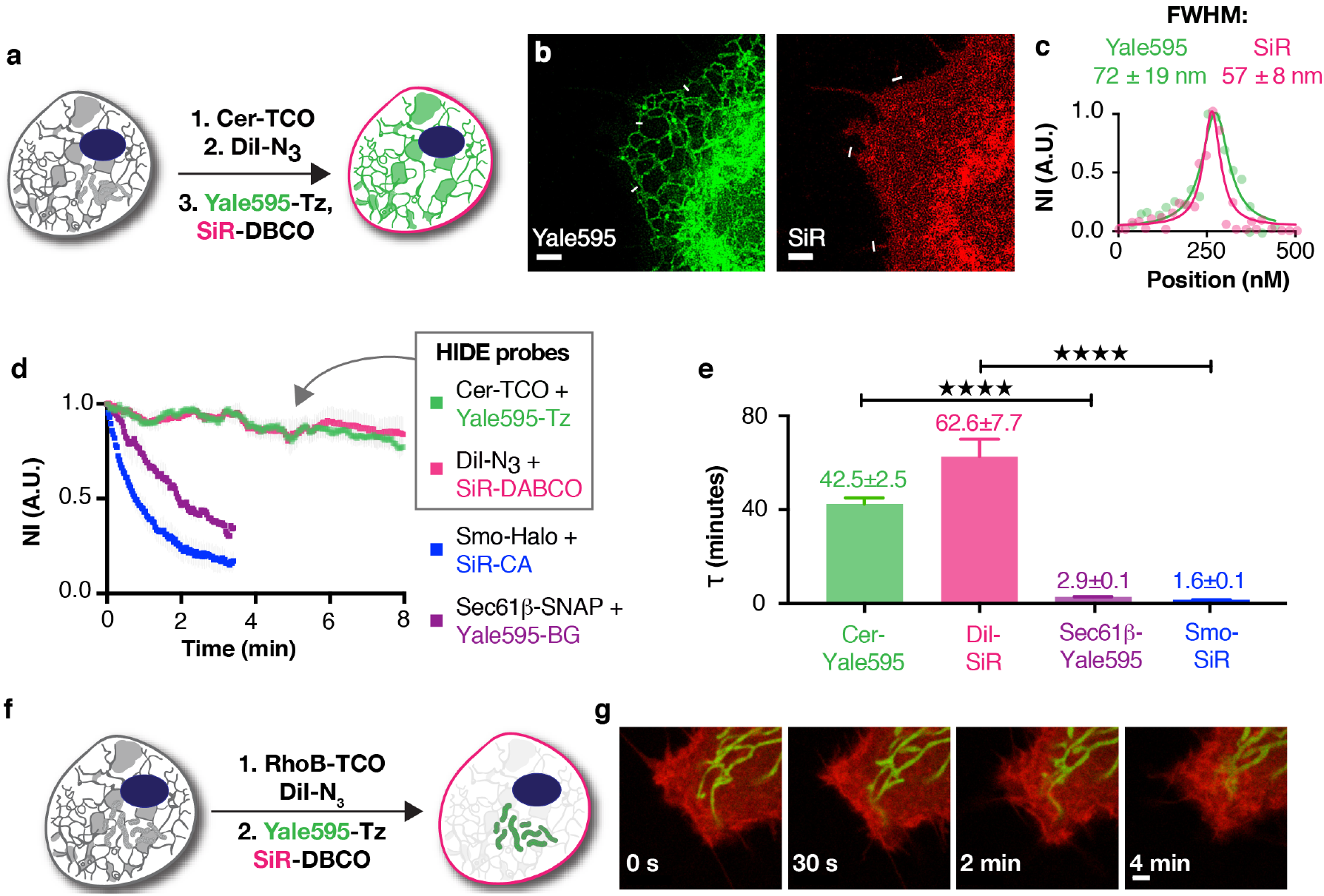
Long time-lapse two-color live-cell STED imaging of HeLa cells. (**a**) Schematic illustration of the three-step procedure employed to label the plasma membrane and ER. (**b**) Two-color STED image of the plasma membrane and ER of HeLa cells. Images from both channels are shown. Scale bars: 2 μM. (**c**) A plot of the fluorescence signal in the two-color STED image as a function of position along the line profile in **b**. A fit of the line profile from the Yale595 channel to a Lorentzian function (green line) provides a FWHM of 72 ± 19 nm. A fit of the line profile from the SiR channel to a Lorentzian function (red line) provides a FWHM of 57 ± 8 nm. (**d**) Plot of normalized fluorescence intensity of protein tags and HIDE probes over time (mean±SD, n = 3 ROI). The fluorescence intensity was measured in each channel separately. (**e**) τ values calculated from a single exponential fit to the photo-bleaching curves in **d** (mean ± SD., n = 3 ROI). ****P ≤ 0.0001. (**f**) Schematic illustration of the two-step procedure employed to label the plasma membrane and mitochondria. (**g**) Time course images of the plasma membrane and mitochondria. Scale bar: 2 μM.

As HIDE probes are generated from pairs of cell-permeant small molecules, they can be used to label both primary and hard-to-transfect cells^6^. To highlight this versatility, we imaged pairs of organelles in two colors by STED in three types of primary cells: human umbilical vein endothelial cells (HUVEC), mouse hippocampal neurons, and retinal pigment epithelium (RPE) cells (Figure 3). Two-color images of the PM and ER of HUVEC cells with SiR-TCO/Yale595-Tz and DiI-N_3_/SiR-DBCO revealed filopodia of one cell strikingly proximal to the ER of an adjacent cell (see ROI I and II in Figure 3b and c, Movie S10). These interactions persisted for several minutes (Figure 3b, arrows). Interestingly while the ER in a single cell is known to form contacts with the PM^20^, the inter-cell interactions evident here have previously not been observed and may represent a new site of inter-cellular communication. In another example, mouse hippocampal neurons were labeled with the dual HIDE PM and mitochondria probes and imaged by STED (Figure 3e, f). We can discern two separate structures, dendritic membrane and mitochondria, only 114 nm apart (Figure 3f, ROI II, Figure 3h). We also observed interactions between dendritic filopodia and mitochondria over a few minutes (Figure 3g, movie S11).

**Figure 3.**
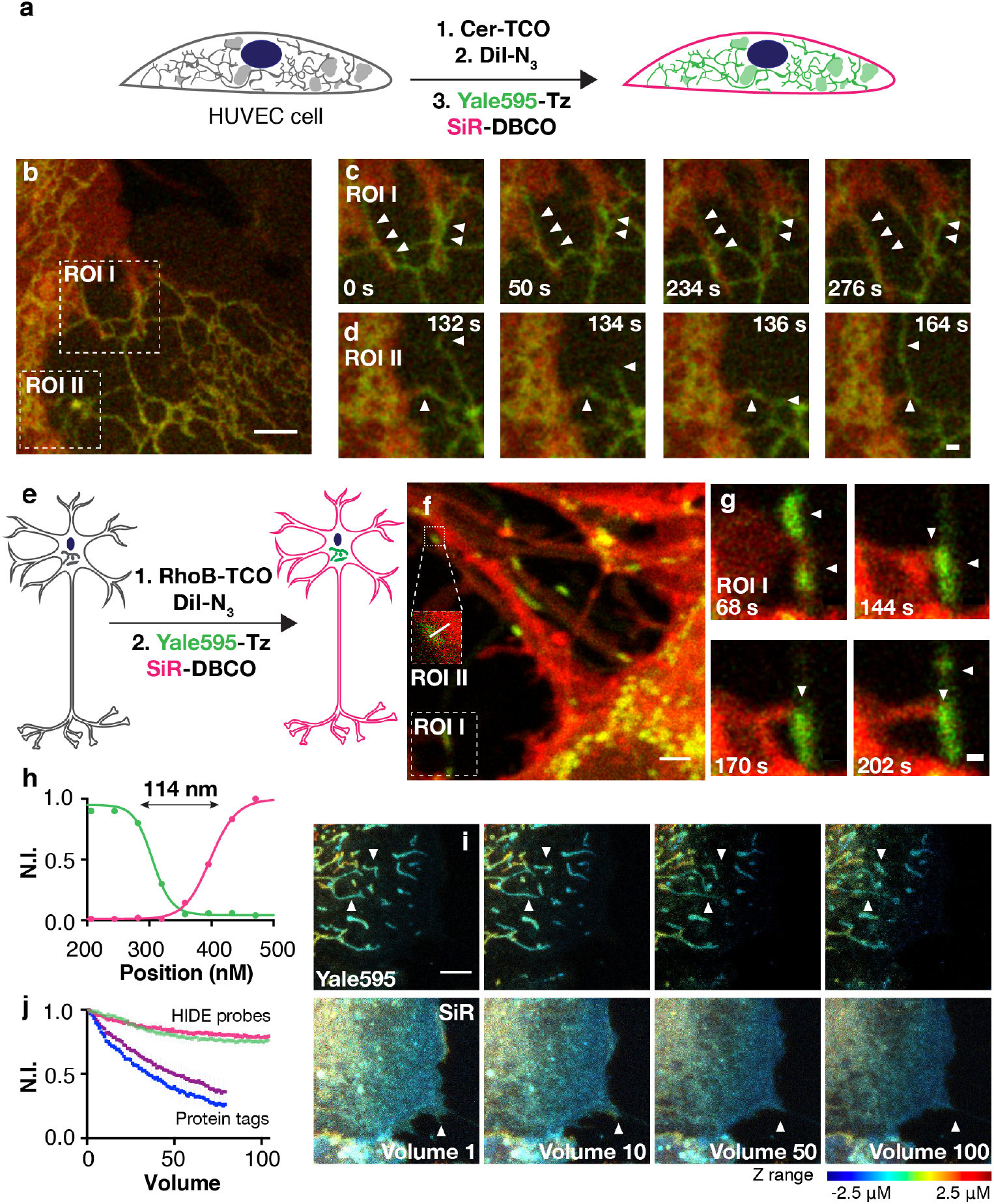
Application of two-color HIDE probes to primary cell lines. (**a**) Schematic illustration of the three-step procedure employed to label the plasma membrane and ER of Human umbilical vein endothelial cells (HUVECs). (**b**) Snapshot of a two-color STED movie of HUVEC. Scale bar: 2 μM. (**c,d**) Time lapse images of ER dynamics and interactions between filopodia and ER. Scale bars: 500nM. (**e**) Schematic illustration of the two-step procedure employed to label the plasma membrane and mitochondria of DIV4 mouse hippocampal neurons. (**f**) Snapshot of a two-color STED movie of DIV4 mouse hippocampal neurons. Scale bar: 2 μM. (**g**) Time lapse images of interactions between filopodia and mitochondria. Scale bars: 500 nM. (**h**) Plot of line profile shown in (**f**, ROI II) illustrating the distance between plasma membrane and mitochondria. (**i**) Time-lapse two-color confocal imaging of mitochondria and plasma membrane in retinal pigment epithelium (RPE) cells. The mitochondrial and plasma membrane volumetric dynamics are recorded continuously over seconds. The axial information is color-coded. 20 z-stacks per volume. volumn rate: 6.1 s. Scale bar: 2 μM (**j**) Plot illustrating normalized fluorescence intensity of RhoB-Yale595 (green), Dil-SiR (red), SM025-Yale595 (purple) and Smo-SiR (blue) over time (mean±SD, n =3 ROI, N = 1 cell).

Importantly the benefits of two-component HIDE probes are not limited to STED. A common photon-demanding imaging modality is 3D time-lapse (4D) imaging. To test the benefit of dual color HIDE probes, we compared conventional confocal 4D imaging of dual HIDE probes to cells tagged with the protein markers SMO25-Yale595 (mitochondria) and Smo-SiR (PM). Here too there was a major benefit of imaging with HIDE probes as SiR and Yale595 tagged HIDE probes were up to 8 times more photostable than the comparable protein-tags (Figure 3i, j).

The new HIDE probes developed here enabled two-color time lapse live-cell STED imaging of more than 250 time points, far exceeding existing examples using state-of-the-art protein tags. The labeling protocols can easily be adapted to various cell lines, including hard-to-transfect and primary cells. We envision that imaging experiments that require a large photon budget would generally benefit from these densely labeled and photostable organelle HIDE probes.

## Acknowledgement

This work was supported by the Wellcome Trust (095927/A/11/Z) and in part by the NIH grants GM 83257 (A.S.), GM 131372 (A.S.), GM 118486 (D.K.T.), NS 089662 (A.J.K.), NS105640 (A.J.K. and Michael J. Higley), MH115939 (A.J.K.), F31MH116571 (J.E.S.), T32GM007223 (J.E.S.) and S10 OD020142 (Leica SP8). L.C. was supported by Brown-Coxe Postdoctoral Fellowship from Yale School of Medicine. We thank Al Mennone (Yale University School of Medicine) for assistance with STED microscopy.

## Author contributions

D.T., A.S. and L.C. designed the experiments. L.C. performed the chemical synthesis and imaging experiments. J.T. measured photophysical properties of dyes. J.E.S. cultured the neuron cells and performed the neuron imaging with L.C. F.R.M. cultured the HUVEC cells. D.T., A.S. and L.C. wrote the manuscript with feedbacks from all other authors.

## Methods

### 1. General methods: chemical synthesis, cell culture, plasmid construction, and microscopy

#### 1.1 Chemical synthesis

Synthesis protocol and additional characterization can be found in the supplementary information.

#### 1.2 Cell culture

HeLa cells (ATCC) were cultured in Dulbecco’s modified Eagle medium (DMEM) (Gibco) supplemented with 10% FBS (Sigma-Aldrich), penicillin (100 unit/mL) and streptomycin (100 μg/mL). hTERT RPE-1 cells (ATCC, CRT-400) were cultured in DMEM/F12 (Gibco) supplemented with 10% FBS, 1% nonessential amino acids (Gibco), 2 mM sodium pyruvate (Gibco), penicillin (100 unit/mL) and streptomycin (100 μg/mL). Human umbilical vein endothelial cells (HUVEC) (Lonza, C2517A) were cultured in EGMTM-2 Endothelial Cell Growth Medium-2 BulletKit (Lonza, CC-3162). All cells were purchased from commercial sources and periodically tested for mycoplasma with DNA methods. Primary neurons were isolated from P0-P1 mouse hippocampus using papain digestion and plated in Neurobasal A media (Invitrogen) with 2% Gem21 supplement (Gemini) and 10% FBS on glass-bottom culture dishes (Matek) coated with poly-D-lysine (20 μg/mL, Corning) and laminin 111 (1 μg/mL, Corning). Media was changed on the neurons after 4 hours and maintained in serum-free Neurobasal A/Gem21 media containing 1% pen/strep and 2 mM L-glutamine.

#### 1.3 Plasmid

The SNAP-Tag plasmid Sec61β-SNAP and OMP25-SNAP was obtained from the Rothman lab and the Bewersdorf lab at Yale School of Medicine^16^.

##### pLVX-ss-HaloTag-Smo

The ss-HaloTag-Smo fragment was PCR amplified from pC4S1-ss-Halotag-mSmo^7^ and cloned by In-Fusion HD into pLVX-puro digested with *EcoRI* and *BamHI* to generate pLVX-ss-pH-Smo for lentivirus production.

#### 1.4 Confocal microscopy

Spinning-disk confocal microscopy was performed using a Improvision UltraVIEW VoX system (Perkin-Elmer) built around a Olympus XI71 inverted microscope, equipped with PlanApo objectives (60×, 1.45 NA) and controlled by the Volocity software (Improvision). To image DiI and SiR, 561-nm and 640-nm laser lines with appropriate filters (615 ± 35 and 705 ± 45 nm, respectively) were used.

#### 1.5 STED microscopy

Live-cell STED microscopy was performed on a Leica TCS SP8 Gated STED 3X microscope. The microscope is equipped with a tunable (460-660 nm) pulsed white light laser for excitation and two HyD detectors for tunable spectral detection. There are also two additional PMT detectors. The microscope is outfitted with three STED depletion lasers (592 nm, 660 nm, and 775 nm). For live cell imaging, the microscope was equipped with a Tokai Hit stage top incubator (model: INUBG2A-GSI) with temperature and CO2 control to maintain an environment of 37°C and 5% CO2. Detailed information on excitation laser and detection window varies depend on the dyes and protocols used and is indicated below in each section. Imaging was conducted with a 100x oil immersion objective (HC PL APO 100x/1.40 OIL) at 1000 Hz with 2 line accumulations in a 19.38 μm2 field of view (1024×1024 pixels at 18.94 nm/pixel). Raw microscopy data were Gaussian blurred (2.0 pixels) in ImageJ. The FWHM values were obtained by fitting line profiles to a Lorentz distribution using Origin 9.1 (www.originlab.com)

### 2. Live-cell imaging of plasma membrane labeled with DiI-N_3_ and SiR-DBCO

#### 2.1 Labeling of plasma membrane with DiI-N_3_ and SiR-DBCO

HeLa cells were incubated with 500 μL of 10 μM DiI-N_3_ in PBS containing 1% casein hydrolysate for 3 min at 37 °C. The cells were then washed and incubated with 500 μL of 2 μM SiR-DBCO in PBS containing 1% casein hydrolysate for 30 min at 37 °C. After washing, DMEM ph(-) was added as imaging buffer, and confocal imaging was performed at 37 °C.

#### 2.2 Optimization of labeling conditions

First, the concentration of DiI-N_3_ was optimized. HeLa cells were incubated with 500 μL of 5 μM, 10 μM, 15 μM, or 20 μM in PBS containing 1% casein hydrolysate DiI-N_3_ for 3 min at 37 °C. After washing, DMEM ph(-) was added as imaging buffer, and confocal imaging was performed at 37 °C. Then the concentration of SiR-DBCO was optimized. HeLa cells were incubated with 500 μL of 10 μM DiI-N_3_ in PBS containing 1% casein hydrolysate for 3 min at 37 °C. The cells were then washed and incubated with 500 μL of 2 μM, 5μM, or 10 μM SiR-DBCO in PBS containing 1% casein hydrolysate for 30 min at 37 °C. After washing, DMEM ph(-) was added as imaging buffer, and confocal imaging was performed at 37 °C.

#### 2.3 Labeling of plasma membrane with pre-mixed DiI-N_3_ and SiR-DBCO

500 μL of 10 μM DiI-N_3_ and 2 μM, 5 μM, or 10 μM SiR-DBCO in PBS containing 1% casein hydrolysate was incubated at 37 °C for 1 hour. The formation of the product was confirmed by MS analysis. The resulting mixture was added to HeLa cells and incubated for 3 min at 37 °C. After washing, DMEM ph(-) was added as imaging buffer, and confocal imaging was performed at 37 °C.

#### 2.4 Measurement of the photostability of DiI-N_3_ and SiR-DBCO

HeLa cells were labeled as described in 2.1 and imaged under STED microscope at 37 °C. SiR dye was excited at 633 nm (40% power) and their emission was detected using a HyD detector from 650-737 nm. The 775 nm depletion laser was used for STED microscopy (30% power). Images were recorded continuously at 2 seconds per frame. Photobleaching plots were generated by normalizing background-subtracted fluorescence intensities (Plasma Membrane ROI – Background ROI) from three separate ROIs using the ROI manager Multi Measure tool in ImageJ.

### 3. Measurement of the photostability of 580CP, JF585, Atto590 and SiR700

HeLa cells were incubated with 500 μL of 10 μM DiI-TCO in PBS containing 1% casein hydrolysate for 3 min at 37 °C. The cells were then washed and incubated with 500 μL of 2 μM 580CP-Tz, JF585-Tz, Atto590-Tz or SiR700 in PBS containing 1% casein hydrolysate for 30 min at 37 °C. After washing, DMEM ph(-)was added as imaging buffer, and STED imaging was performed at 37 °C. 580CP, and Atto590 were excited at 590 nm (5% power) and their emission was detected using a HyD detector from 600-670 nm. JF585 dye was excited at 590 nm (15% power) and their emission was detected using a HyD detector from 600-670 nm. SiR700 dye was excited at 650 nm (40% power) and their emission was detected using a HyD detector from 660-730 nm. The 775 nm depletion laser was used for STED microscopy (30% power). Images were recorded continuously at 2 seconds per frame. Photobleaching plots were generated by normalizing background-subtracted fluorescence intensities (Plasma Membrane ROI – Background ROI) from three separate ROIs using the ROI manager Multi Measure tool in ImageJ.

### 4. Live-cell imaging with Yale595

#### 4.1 Measurement of photophysical properties of Yale595

##### 4.1.1 Absorption and Emission Spectra of Yale-595 and SiR

The absorbance spectra of Yale-595 and SiR were measured at 2 μM in DPBS (w/ 0.1% DMSO) on a Beckman UV-Vis spectrophotomerter. The emission spectra of the same solutions were measured on a TCSPC TD-Fluor Horiba Fluorolog 3 Time Domain Fluorimeter.

##### 4.1.2 Quantum Yield Determination for Yale-595

Solutions of Yale-595 and Bodipy-Texas Red between 0 – 10 μM were prepared in DPBS (w/ 0.05% - 0.25% DMSO). Absorbance at 550 nm was measured for each on a Beckman UV-Vis spectrophotomerter. Afterwards, fluorescence of the same samples was measured on a TCSPC TD-Fluor Horiba Fluorolog 3 Time Domain Fluorimeter. Both the excitation and emission bandwidths for all measurements were set to 5 nm. The spectra were recorded with a step size of 1 nm. The excitation wavelength was set to 550 nm and the emission spectra were recorded in the 560-800 nm interval. Absorbance at 550 nm versus integrated fluorescence intensity for each solution was plotted using GraphPad Prism 7.0. The relative quantum yield of Yale-595 was calculated from the linear regressions according to the following:

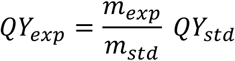

Where *QY*_*exp*_ and *m*_*exp*_ are the relative quantum yield and slope of linear regression, respectively, for Yale-595 and *QY*_*std*_ and *m*_*std*_ are the absolute quantum yield and slope of linear regression, respectively, for Bodipy-Texas Red. The absolute quantum yield for Bodipy-Texas Red was previously reported.

##### 4.1.3 Standard Curve of Exctinction Coefficient for Yale 595

Solutions of Yale-595 and SiR at 5μM, 10 μM, 15 μM, 20 μM and 25 μM in DPBS (w/ 0.25 – 1.25% DMSO) were prepared. The absorbance at 597 nm (Yale-595) or 652 nm (SiR) were measured on Spiegel Lab plate reader using polypropylene 96-well plates. The concentrations were plotted against their absorbances using GraphPad Prism 7.0. The extinction coefficient of Yale-595 was calculated from the linear regressions according to the following:

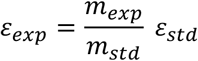

Where *ε*_*exp*_ and *m*_*exp*_ are the extinction coefficient and slope of linear regression, respectively, for Yale-595 and *ε*_*std*_ and *m*_*std*_ are the extinction coefficient and slope of linear regression, respectively, for SiR. The extinction coefficient for SiR was previously reported^8^.

#### 4.2 Measurement of absorbance in response to different solvent polarities

Yale595-COOH (or JF585-COOH) was dissolved into a mixture of water and dioxane (0-100% dioxane in water) to a final concentration of 10 μM. The absorbance data was recorded on a DU730 Life Science UV/Vis spectrophotometer using the wavelength scan mode.

#### 4.3 Measurement of the photostability of Yale595

HeLa cells were incubated with 500 μL of 10 μM DiI-TCO in PBS containing 1% casein hydrolysate for 3 min at 37 °C. The cells were then washed and incubated with 500 μL of 2 μM Yale595-Tz in PBS containing 1% casein hydrolysate for 30 min at 37 °C. After washing, DMEM ph(-)was added as imaging buffer, and STED imaging was performed at 37 °C. Yale595 dye was excited at 595 nm (10% power) and their emission was detected using a HyD detector from 605-675 nm. The 775 nm depletion laser was used for STED microscopy (30% power). Images were recorded continuously at 2 seconds per frame. Photobleaching plots were generated by normalizing background-subtracted fluorescence intensities (Plasma Membrane ROI – Background ROI) from three separate ROIs using the ROI manager Multi Measure tool in ImageJ.

#### 4.4 Live-cell STED imaging of ER using Yale595

HeLa cells were incubated with 500 μL of 2 μM Cer-TCO in PBS containing 1% casein hydrolysate and 0.2% Pluronic F-127 for 60 min at 37 °C. The cells were then washed and incubated with 500 μL of 2 μM Yale595-Tz in PBS containing 1% casein hydrolysate for 30 min at 37 °C and imaged on the SP8 STED microscope. Yale595 dye was excited at 595 nm (10% power) and their emission was detected using a HyD detector from 605-675 nm. The 775 nm depletion laser was used for STED microscopy (30% power).

### 5. Live-cell two-color STED imaging using HIDE probes

#### 5.1 Plasma membrane and ER of HeLa cells

HeLa cells were incubated with 500 μL of 4 μM Cer-TCO in Live Cell Imaging Solution (Life Technologies, A14291DJ) containing 0.2% Pluronic F-127 for 60 min at 37 °C. After washing, the cells were treated with 500 μL of 10 μM Dil-N_3_ in Live Cell Imaging Solution for 3 min at 37 °C. After washing, the cells were incubated with 500 μL of 2 μM SiR-DBCO and 2 μM Yale595-Tz in Live Cell Imaging Solution for 30 min at 37 °C. After washing, STED imaging was carried out at 37 °C using DMEM ph(-) as the imaging buffer under Leica SP8 microscope with two excitation lasers at 595 nm (10% power), 650 nm (60% power) and two detection windows at 605-625 nm and 670-750 nm. The 775 nm depletion laser was used for STED microscopy (30% power). Images were recorded continuously at 2 seconds per frame. Raw image data were spectralunmixed and Gaussian blurred (2.0 pixels) in ImageJ.

#### 5.2 Plasma membrane and mitochondria of HeLa cells

HeLa cells were incubated with 500 μL of 10 μM RhoB-TCO and 10 μM Dil-N_3_ in Live Cell Imaging Solution for 3 min at 37 °C. After washing, the cells were incubated with 500 μL of 2 μM SiR-DBCO and 2 μM Yale595-Tz in Live Cell Imaging Solution for 30 min at 37 °C. After washing, cells were incubated at 37 °C in DMEM ph(-) for 30 min to wash out the remaining dyes. After further washing, STED imaging was carried out at 37 °C using DMEM ph(-) as the imaging buffer under Leica SP8 microscope with two excitation lasers at 595 nm (10% power), 650 nm (60% power) and two detection windows at 605-625 nm and 670-750 nm. The 775 nm depletion laser was used for STED microscopy (30% power). Images were recorded continuously at 2 seconds per frame. Raw image data were spectral unmixed and Gaussian blurred (2.0 pixels) in ImageJ.

#### 5.3 Plasma membrane and ER of Human umbilical vein endothelial cells (HUVEC)

HUVECs were incubated with 500 μL of 4 μM Cer-TCO in Live Cell Imaging Solution containing 0.2% Pluronic F-127 for 60 min at 37 °C. After washing, the cells were treated with 500 μL of 10 μM Dil-N_3_ in Live Cell Imaging Solution for 3 min at 37 °C. After washing, the cells were incubated with 500 μL of 2 μM SiR-DBCO and 2 μM Yale595-Tz in Live Cell Imaging Solution for 30 min at 37 °C. After washing, STED imaging was carried out at 37 °C using DMEM ph(-) as the imaging buffer under Leica SP8 microscope with two excitation lasers at 594 nm (10% power), 650 nm (60% power) and two detection windows at 605-625 nm and 670-750 nm. The 775 nm depletion laser was used for STED microscopy (30% power). Images were recorded continuously at 2 seconds per frame. Raw image data were spectral unmixed and Gaussian blurred (2.0 pixels) in ImageJ.

#### 5.4 Plasma membrane and mitochondria of mouse hippocampal neuron cells

Neuron cells were incubated with 500 μL of 10 μM RhoB-TCO and 10 μM Dil-N_3_ in DMEM for 3 min at 37 °C. After washing, the cells were incubated with 500 μL of 2 μM SiR-DBCO and 2 μM Yale595-Tz in DMEM for 30 min at 37 °C. After washing, cells were incubated at 37 °C in DMEM ph(-) for 30 min to wash out the remaining dyes. After further washing, STED imaging was carried out at 37 °C using DMEM as the imaging buffer under Leica SP8 microscope with two excitation lasers at 595 nm (10% power), 650 nm (60% power) and two detection windows at 605-625 nm and 670-750 nm. The 775 nm depletion laser was used for STED microscopy (30% power). Images were recorded continuously at 2 seconds per frame. Raw image data were Gaussian blurred (2.0 pixels) in ImageJ.

### 6. Live-cell two-color imaging using protein tags

#### 6.1 Two-color STED imaging of plasma membrane and ER

Generation of hTERT RPE-1 cells stably expressing Smo-Halo was described previously using pLVX-ss-HaloTag-Smo^21^. hTERT RPE-1 cells stably expressing Smo-Halo were transfected with Sec61β-SNAP using FuGENE HD Transfection Reagent (Promega) one day before imaging. Before imaging, the cells were incubated with 500 μL of 2 μM SiR-CA and 2 μM Yale595-BG in Live Cell Imaging Buffer for 30 min at 37 °C. After washing, STED imaging was carried out at 37 °C using DMEM as the imaging buffer under Leica SP8 microscope with two excitation lasers at 595 nm (10% power), 650 nm (60% power) and two detection windows at 605-625 nm and 670-750 nm. The 775 nm depletion laser was used for STED microscopy (30% power). Images were recorded continuously at 2 seconds per frame. Raw image data were spectral unmixed Gaussian blurred (2.0 pixels) in ImageJ.

#### 6.2 Two-color confocal 3D imaging of plasma membrane and Mitochondria

hTERT RPE-1 cells stably expressing Smo-Halo were transfected with OMP25-SNAP using FuGENE HD Transfection Reagent (Promega) one day before imaging. Before imaging, the cells were incubated with 500 μL of 2 μM SiR-CA and 2 μM Yale595-BG in Live Cell Imaging Buffer for 30 min at 37 °C. After washing, STED imaging was carried out at 37 °C using DMEM as the imaging buffer under Leica SP8 microscope with two excitation lasers at 595 nm (10% power), 650 nm (60% power) and two detection windows at 605-625 nm and 670-750 nm. The 775 nm depletion laser was used for STED microscopy (30% power). Images were recorded continuously at 2 seconds per frame. Raw image data were spectral unmixed Gaussian blurred (2.0 pixels) in ImageJ.

### 7. Live-cell two-color 3D confocal imaging of plasma membrane and mitochondria using HIDE probes

HeLa cells were incubated with 500 μL of 10 μM RhoB-TCO and 10 μM Dil-N_3_ in Live Cell Imaging Solution for 3 min at 37 °C. After washing, the cells were incubated with 500 μL of 2 μM SiR-DBCO and 2 μM Yale595-Tz in Live Cell Imaging Solution for 30 min at 37 °C. After washing, cells were incubated at 37 °C in DMEM ph(-) for 30 min to wash out the remaining dyes. After further washing, 3D confocal imaging was carried out at 37 °C using DMEM ph(-) as the imaging buffer under Leica SP8 microscope with two excitation lasers at 595 nm (3% power), 650 nm (20% power) and two detection windows at 605-625 nm and 670-750 nm. The images were taken at 20 z-stacks per volume at 6.1s per volume. Raw image data were spectral unmixed in ImageJ.

### 8. Evaluation of cellular toxicity of two-color HIDE probes using a cell division assay^6^

We monitored the cell division in three groups of HeLa cells. 1) Non-treated cells; 2) HeLa cells labeled as described in 5.1; and 3) HeLa cells labeled as described in 5.2. Labeling of the cells was confirmed by a confocal microscopy. Wide field imaging was carried out on an Invitrogen™ EVOS™ FL Auto 2 Imaging system equipped with temperature, humidity, and atmosphere controls for live cell imaging. The sample was kept at 37 °C with 5% CO2 and imaged using a 20x objective and a Highsensitivity 1.3 MP CMOS monochrome camera (1,328 × 1,048 pixels) every 10 minutes for 15 h using bright field. Cell division events were counted manually.

## References

1. Abbe, E. Beiträge zur Archiv. Mikrosk. Anat. 9, 413–418 (1873).

2. Sahl, S.J., Hell, S.W. & Jakobs, S. Nat Rev Mol Cell Bio 18, 685–701 (2017).

3. Nixon-Abell, J. et al. Science 354 (2016).

4. Oracz, J., Westphal, V., Radzewicz, C., Sahl, S.J. & Hell, S.W. Sci Rep-Uk 7 (2017).

5. Erdmann, R.S. et al. Angew Chem Int Edit 53, 10242–10246 (2014).

6. Thompson, A.D. et al. Angew Chem Int Edit 56, 10408–10412 (2017).

7. Takakura, H. et al. Nat Biotechnol 35, 773–780 (2017).

8. Lukinavicius, G. et al. Nat Chem 5, 132–139 (2013).

9. Blackman, M.L., Royzen, M. & Fox, J.M. J Am Chem Soc 130, 13518–13519 (2008).

10. Devaraj, N.K., Weissleder, R. & Hilderbrand, S.A. Bioconjugate Chem 19, 2297–2299 (2008).

11. Han, Y.B., Li, M.H., Qiu, F.W., Zhang, M. & Zhang, Y.H. Nat Commun 8 (2017).

12. Chen, B.C. et al. Science 346, 1257998 (2014).

13. Agard, N.J., Prescher, J.A. & Bertozzi, C.R. J Am Chem Soc 127, 11196–11196 (2005).

14. Debets, M.F. et al. Chem Commun 46, 97–99 (2010).

15. Butkevich, A.N. et al. Angew Chem Int Edit 55, 3290–3294 (2016).

16. Bottanelli, F. et al. Nat Commun 7 (2016).

17. Neumann, D., Buckers, J., Kastrup, L., Hell, S.W. & Jakobs, S. PMC Biophys 3, 4 (2010).

18. Grimm, J.B. et al. Nat Methods 14, 987–994 (2017).

19. Lukinavicius, G. et al. J Am Chem Soc 138, 9365–9368 (2016).

20. Phillips, M.J. & Voeltz, G.K. Nat Rev Mol Cell Bio 17, 69–82 (2016).

21. Kukic, I., Rivera-Molina, F. & Toomre, D. Cilia 5, 23 (2016).

